# MyROOT: A novel method and software for the semi-automatic measurement of plant root length

**DOI:** 10.1101/309773

**Authors:** Isabel Betegón-Putze, Alejandro González, Xavier Sevillano, David Blasco-Escámez, Ana I. Canño-Delgado

## Abstract

Root analysis is essential for both academic and agricultural research. Despite the great advances in root phenotyping and imaging however, calculating root length is still performed manually and involves considerable amounts of labor and time. To overcome these limitations, we have developed MyROOT, a novel software for the semi-automatic quantification of root growth of seedlings growing directly in agar plates. Our method automatically determines the scale from the image of the plate, and subsequently measures the root length of the individual plants. To this aim, MyROOT combines a bottom-up root tracking approach with a hypocotyl detection algorithm. At the same time as providing accurate root measurements, MyROOT also significantly minimizes the user intervention required during the process. Using Arabidopsis, we tested MyROOT with seedlings from different growth stages. Upon comparing the data obtained using this software with that of manual root measurements, we found that there are no significant differences (t-test, p-value < 0.05). Thus, MyROOT will be of great aid to the plant science community by permitting high-throughput root length measurements while saving on both labor and time.

## INTRODUCTION

The root, which is responsible for anchoring the plant to the soil, is an essential organ for overall plant growth and development. The characterization of different root traits is therefore important not only for understanding organ growth, but also for evaluating the impact of roots in agriculture ^1^. As such, generating tools for precise, high-throughput phenotyping and imaging of the root is essential for plant research and agriculture. Even phenotyping facilities such as the ones available in the European Plant Phenotypic Network (http://www.plant-phenotyping-network.eu/) have started to implement tools for the massive screening of roots.

Roots provide the necessary structural and functional support for the incorporation of nutrients and water from the soil. In *Arabidopsis thaliana* (Arabidopsis), the primary root has a very simplified anatomy that makes it very amenable for genetic and microscopic analyses ^2–4^ Different root cell lineages are derived from the activity of a group of stem cells located at the root apex. Here, the stem cell niche is formed by a few (3-7) quiescent center (QC) cells that occasionally divide asymmetrically to renew themselves and to form daughter stem cells. From the root apex, these cells actively divide in the meristematic zone, and before exiting the cell cycle in the transition zone, continue to elongate and differentiate in spatially separated regions of the root. In this way, primary root growth is determined by the balance between cell division and cell elongation within the different zones of the root ^5–8^.

The most straightforward symptom of abnormal root growth or development can be identified by examining the length of the primary root in seedlings. Abnormalities in length can usually be observed and measured just five to six days after germination (DAG), where still reflect their embryonic origin ^9^. Growth defects in the primary root of seedlings are not only consistent with overall growth defects, but also persistent along the entire plant life cycle ^10–12^ Indeed, Arabidopsis root analyses were the foundations for multiple genetic screens that ultimately led to the identification of several key regulators of plant growth and development^10,13–16^.

Root analysis of young seedlings offers direct information regarding overall plant growth and viability. Despite important advances in plant imaging techniques such as microscopic visualization ^17-19^, the root length of seedlings growing in agar plates is generally measured by manually indicating the position of each seedling or manually tracking each root using the ImageJ software (https://imagej.nih.gov/ij/). For this reason, the development and use of methods that enable the automatic analysis of a large number of roots represents a step forward for high-throughput root analysis. Automatic analysis of root system architecture is just beginning to be implemented, and novel methods based on acquiring, processing, and obtaining quantitative data from root images are now available (for a review, see^19^).

With this in mind, our rational was to develop MyROOT, a software capable of semi-automatically calculating root length. By precisely detecting all individual roots and hypocotyls growing on an agar plate from a JPEG image, this software simplifies and minimizes user intervention during the calculation of root length. As a proof of concept, MyROOT software was first used for root length measurement of wild type and brassinosteroid-signaling Arabidopsis mutants grown in control and osmotic stress conditions. MyROOT software is available at https://www.cragenomica.es/research-groups/brassinosteroid-signaling-in-plant-development.

Currently, a myriad of software is available for measuring root phenotypic traits. These differ with respect to: i) the medium in which the plant is grown, ii) the use of 2D or 3D imaging, iii) the imaging modality, and iv) the degree of manual intervention required from the user ^1^. Specifically, when using these available software tools to measure root length, plant scientists face three main hurdles: i) the constraints imposed during the image acquisition process, ii) the ease of use (often related to the degree of manual intervention required), and iii) the accuracy of the root length measurements.

Regarding the complexity of the image acquisition process, software tools such as PlaRoM ^20^ require scanning the plates using a camera-microscope unit mounted on a robotic arm, BRAT ^21^ requires a cluster of several flatbed scanners, and the RootReader2D software ^22^ requires a camera equipped with two cross-polarized filters. In comparison, MyROOT operates on photographs (of Petri dishes) taken directly from above with a standard digital camera or even a good quality cell phone.

In terms of manual intervention, some of the software tools require intensive user participation in order to define certain aspects of the individual roots under analysis. This makes measuring root length a time-consuming and burdensome process. A prime example of this is RootTrace ^23^, a software for which the user has to manually define the start point of each root. In contrast, MyROOT merely requires the user to define the region in which the seedlings are placed on the plate, and then subsequently operates in a fully automatic fashion. Moreover, the interactive interface of MyROOT permits the visualization of results from each intermediate step of the root measurement process. In this way, the user can modify any configuration parameter at will, and redo any of the steps if deemed necessary.

Finally, as far as measurement accuracy is concerned, the precise detection of root start and end points is critical. While some software tools rely on intensive manual labor for defining the aforementioned points ^24^, others such as BRAT ^21^ use shoot detection to determine the root start point. In contrast, to ensure that roots are measured correctly from their tip to their true start point, MyROOT combines a bottom-up root tracking approach with a hypocotyl detection algorithm to provide accurate measurements.

In summary, not only is MyROOT very straightforward to use, but it can also accurately measure root length on a plate while sparing important and unnecessary human labor.

## RESULTS

### MyROOT is a software for the high-throughput analysis of root length

The majority of root studies begin with an overall determination of root growth as estimated by manual, laborious and time-consuming measurements. To address this limitation, we developed a semi-automatic and non-invasive software for the high-throughput measurement of root length. This method is implemented in Matlab as an automatic tool named MyROOT (Fig. 1a). It is based on pictures of whole agar plates where young seedlings are growing vertically on the surface, and implements novel algorithms capable of separately detecting the root and the hypocotyl of each individual seedling and estimating a hypocotyl curve based on the detection of some hypocotyls (Fig. 1b-g).

**Figure 1.**
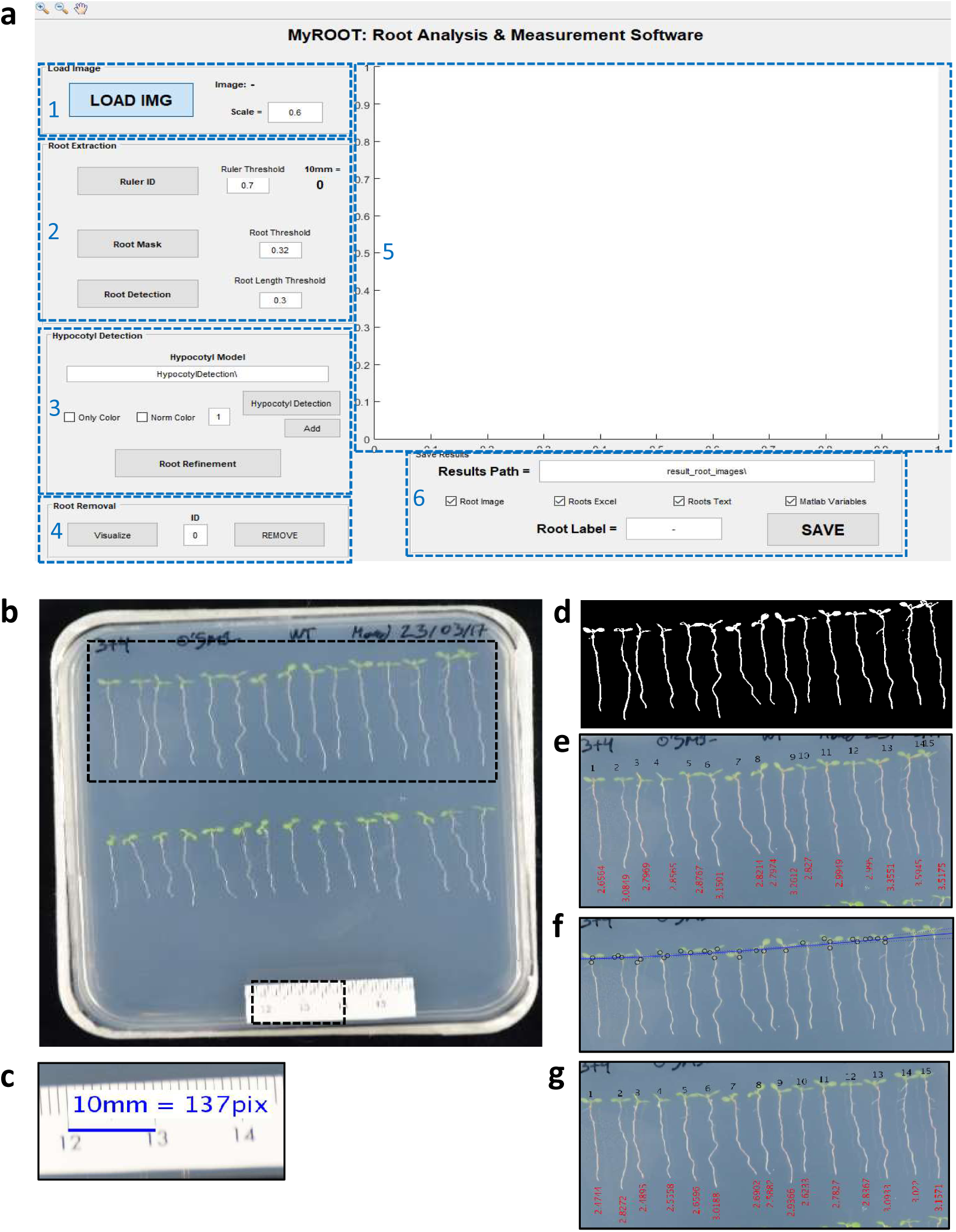
The graphical interface and steps of MyROOT. (**a**) The graphical user interface of MyROOT is organized into six sections: 1. Input image information, 2. Root extraction parameters, 3. Hypocotyl detection parameters, 4. Manual removal of roots, 5. Visualization of the image and the different detection steps, and 6. Saving parameters. (**b**) The input image required for analysis is a picture of the square plate in which the aligned seedlings are growing. By using information from this image, MyROOT performs the following steps: (**c**) Identification of the ruler to determine the scale (i.e., the equivalence between pixels and millimeters), (**d**) Root segmentation to identify the seedlings, (**e**) Root tracking to measure the roots, (**f**) Hypocotyl detection to identify the hypocotyls and separate them from the roots, and (**g**) Root measurement to quantify the length of individual seedlings (i.e., the distance from the root tip to the end of the hypocotyl).

MyROOT detects and measures root length by following a series of steps (Fig. 1b-g). First, a digital image of the plate containing the growing seedlings is taken and used for the analysis (Fig. 1b and Fig. S1). The image is taken with a ruler (at least 1 cm long) placed on top of the plate. From each JPEG image, the software: i) detects 1 cm of the ruler to automatically compute the scale and calculate the equivalence between pixels and millimeters (mm; Fig. 1c); ii) generates a binary mask from the manually selected area that allows for root segmentation (this separates those pixels that belong to a root from those of the background) (Fig. 1d); iii) measures the length of the roots through a root tracking process (Fig. 1e); iv) computes a regression curve based on the detection of the hypocotyls to identify the starting point of each root (Fig. 1f); v) measures the root length again from the root tip to the end of the hypocotyl (Fig. 1g); and vi) exports the measurements and the generated masks to a new folder. Finally, the results are saved in: i) an Excel spreadsheet or a TXT file in which each root is identified by an ID tag, length value and a descriptive text label introduced by the user; ii) an image showing the detected and measured roots; and iii) MATLAB variables including the intermediate data such as hypocotyl position and the detection curve that were generated while quantifying root length.

One of the advantages of this software is that it allows the user to supervise the different steps of the process as the results of each step are displayed before executing the following one. This feature enables the user to modify the different parameters (e.g., segmentation thresholds for ruler and root detection, and model for hypocotyl detection, etc.) at any point in the process to take into account different image conditions. Nonetheless, default parameter values have been set for satisfactory operation on a wide range of images for pre-defined acquisition conditions (see Material and Methods). Furthermore, the position of any hypocotyl that is not automatically detected can be manually indicated, and undesired roots can be manually removed from the results before saving.

In summary, by determining the pixel-millimeter equivalence and detecting seedling morphology (roots and hypocotyls) from an image of a seedling-containing agar plate, MyROOT offers a valuable analytical tool for precisely measuring root growth in a semi-automatic manner. As such, this software clearly provides a solution to the timely task of manually quantifying root length.

### Root detection and measurement process

MyROOT has been developed for the high-throughput, accurate, and non-invasive measurement of root length from seedlings growing in agar plates. In this respect, the three most crucial steps are to precisely determine the scale, identify the roots, and measure their length. The scale information is obtained from a piece of measuring tape that is placed on the surface of the Petri dish. This allows the measurements to be completely independent from the specific characteristics of the image capture system. The first step for detecting the ruler is based on its color contrast with the background. By computing the vertical and horizontal profiles of the image, the algorithm is designed to explore the entire image in search of a white patch (Fig. 2a). As the border of the plate has a similar color contrast with the background, a median filter is applied to reduce the border effect. The maximum values in the filtered profiles define the image area where the white patch is present. Next, the resulting area is further cropped (Fig. 2b) and processed (Fig. 2c-e). By applying a threshold based on Otsu’s algorithm ^25^, the black lines representing cm and mm marks are not filtered out (Fig. 2b). Finally, a horizontal profile of this binary image is generated (Fig. 2d) in which the pixel-mm equivalence is defined as the difference between consecutive local maxima (Fig. 2e).

**Figure 2.**
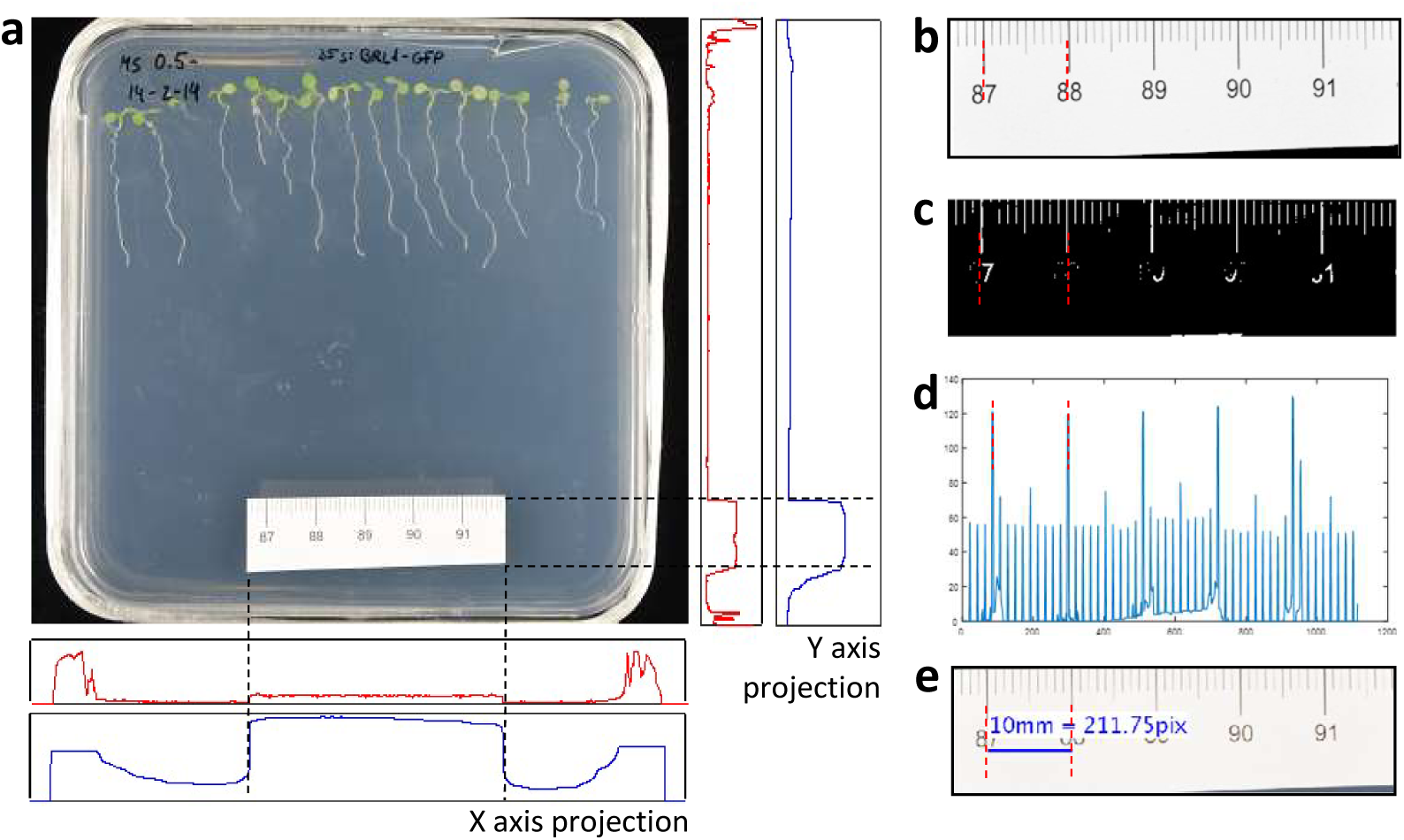
The ruler identification process. (**a**) MyROOT computes the vertical and horizontal profiles of the image to look for a white patch. (**b**) The ruler is identified. (**c**) The area corresponding to the ruler is then segmented into light and dark areas (binarization), for which black lines (dark areas) are identified with high values and white areas with lower values. (**d**) A profile is generated in which black lines are identified as peaks. (**e**) By using the distance between peaks, the equivalence between pixels and millimeters is calculated.

The core of the whole method is the root extraction and measurement process. In order to extract roots, the user must first manually define the area in which roots are present (note: only one row of seedlings should be included when defining the area). Then, with just a few mouse clicks from the user, a binary mask is generated that allows root segmentation. This later leads to the identification of individual roots through a root tracking process, and finally allows the individually identified roots to be measured (Fig. 3). The root segmentation process can be divided into four main steps: i) color normalization (Fig. 3a), ii) ridge detection (Fig. 3b), iii) root tracking (Fig. 3c), and iv) root identification (Fig. 3d). During the color normalization step, the image is processed and a global working framework is set (i.e., all images going through this process become color-balanced and have the same lower and higher white values; Fig. 3a). This allows the user to manage different initial conditions (illumination, color, and saturation, etc.) while continuing with the same subsequent steps of the pipeline. In the next step, a ridge (i.e., white contrasted area) detector identifies roots based on their contrast with the background (for this, the level of whiteness is irrelevant; Fig. 3b). After the detection step, a final mask is generated for tracking the roots. Due to the linear disposition of the roots in the plate, we employed a bottom-up tracking approach. As such, tracking starts at the end point of each root and continues upward, row by row, until the hypocotyl detection curve is found (Fig. 3c). Finally, the tracking of each root makes it possible to identify which pixels correspond to which root (Fig. 3d).

**Figure 3.**
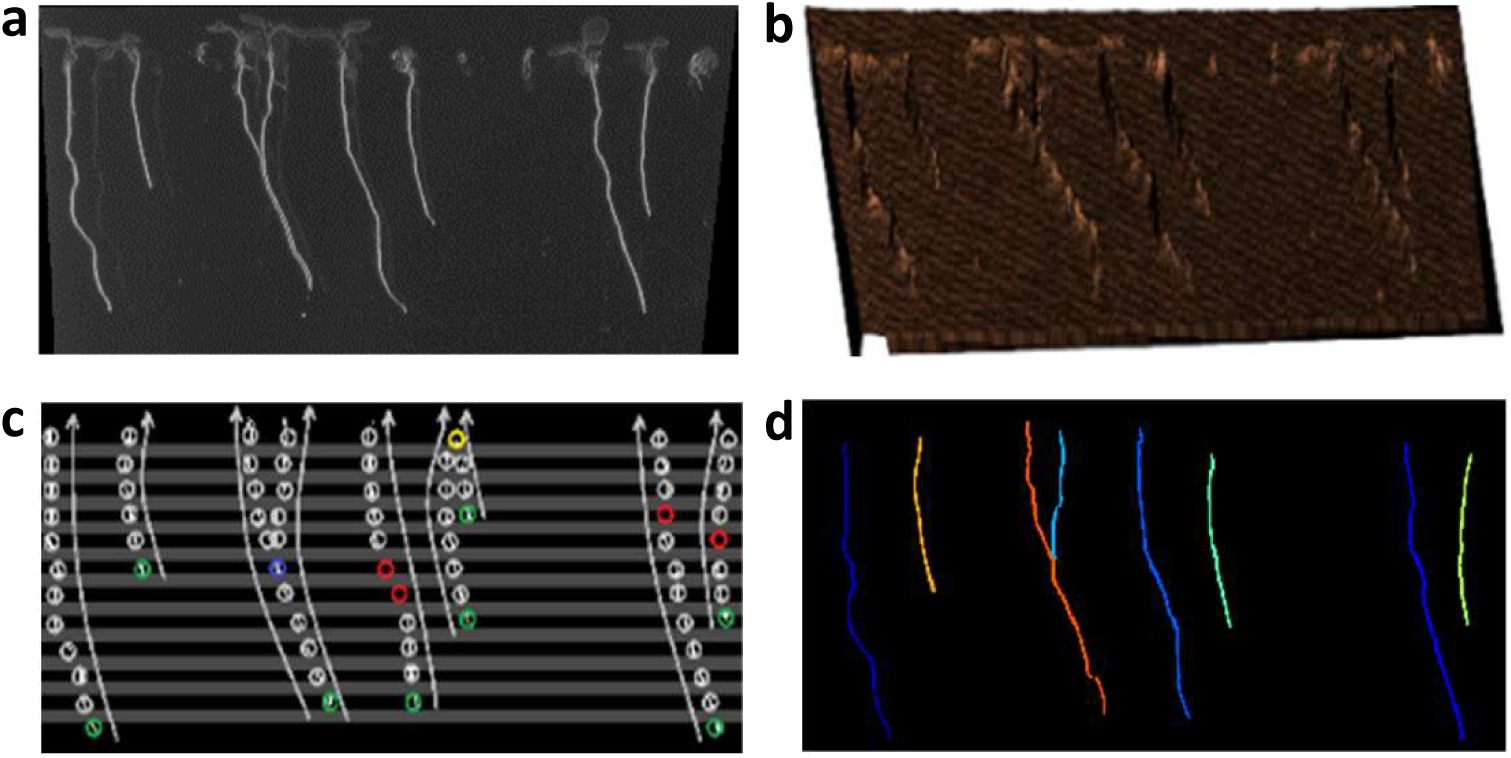
Root extraction method. (**a**) Colors are normalized in the area where roots are present, and white roots are detected. (**b**) Segmentation is performed by applying a ridge detector. (**c**) Starting at the root tip, the roots are tracked using a bottom-up approach. (**d**) Each root is measured using its historical recorded tracking, and root length is calculated by taking into account the pixel-millimeter equivalence.

Once the root tracking process has been completed, each individual root is measured based on previous positions saved in the historical record. Specifically, root length is calculated in pixels by adding the distances between previous consecutive points and then applying the previously calculated pixel-mm equivalence. Next, a refinement process is applied in which very short roots, which are often associated with noise, are discarded. By default, MyROOT discards any root measurement shorter than 30% of the longest one. However, this percentage can be manually chosen by the user if need be. A second filter is then applied in order to keep those roots that terminate close to the previously calculated hypocotyl curve. If a root surpasses the hypocotyl curve, it is cut at this level. Finally, a unique numeric identifier (ID) is assigned to all roots that are not filtered out during processing.

As two roots can be located so close to one another that they cannot be detected as individual roots, we trained MyROOT with the following characteristics: i) when a split occurs and a current root matches more than one detection (blue circle in Fig. 3C), a new root sharing the same historical record is created, and ii) when a fusion occurs and two roots match a single detection (yellow circle in Fig. 3c), the shortest root is eliminated from the root set and added as a sub-root of the longest one.

To validate our software, we compared root length measurements obtained using MyROOT with manual measurements performed using ImageJ. As a first comparison, six-day-old seedlings grown in three different vertical Petri dishes (n=89) were measured by three different scientists (Fig. 4a). Upon comparing these MyROOT and ImageJ measurements, we observed no significant difference (t-test, p-value < 0.05) in the mean root lengths (23.45, 23.61 and 23.62 mm using ImageJ, and 23.47, 23.51 and 23.35 mm using MyROOT). Taken together, these results indicate that measurements made using our software coincide with manual measurements, thereby supporting the use of MyROOT for root length analysis. Furthermore, a second comparison was conducted by measuring the root length of the same seedlings during six consecutive days (from three to eight DAG; n>116; Fig. 4b). Again, highly similar values were obtained between the MyROOT and manual measurements. Importantly, this validates MyROOT for analyzing the root length of Arabidopsis seedlings at different days after germination.

**Figure 4.**
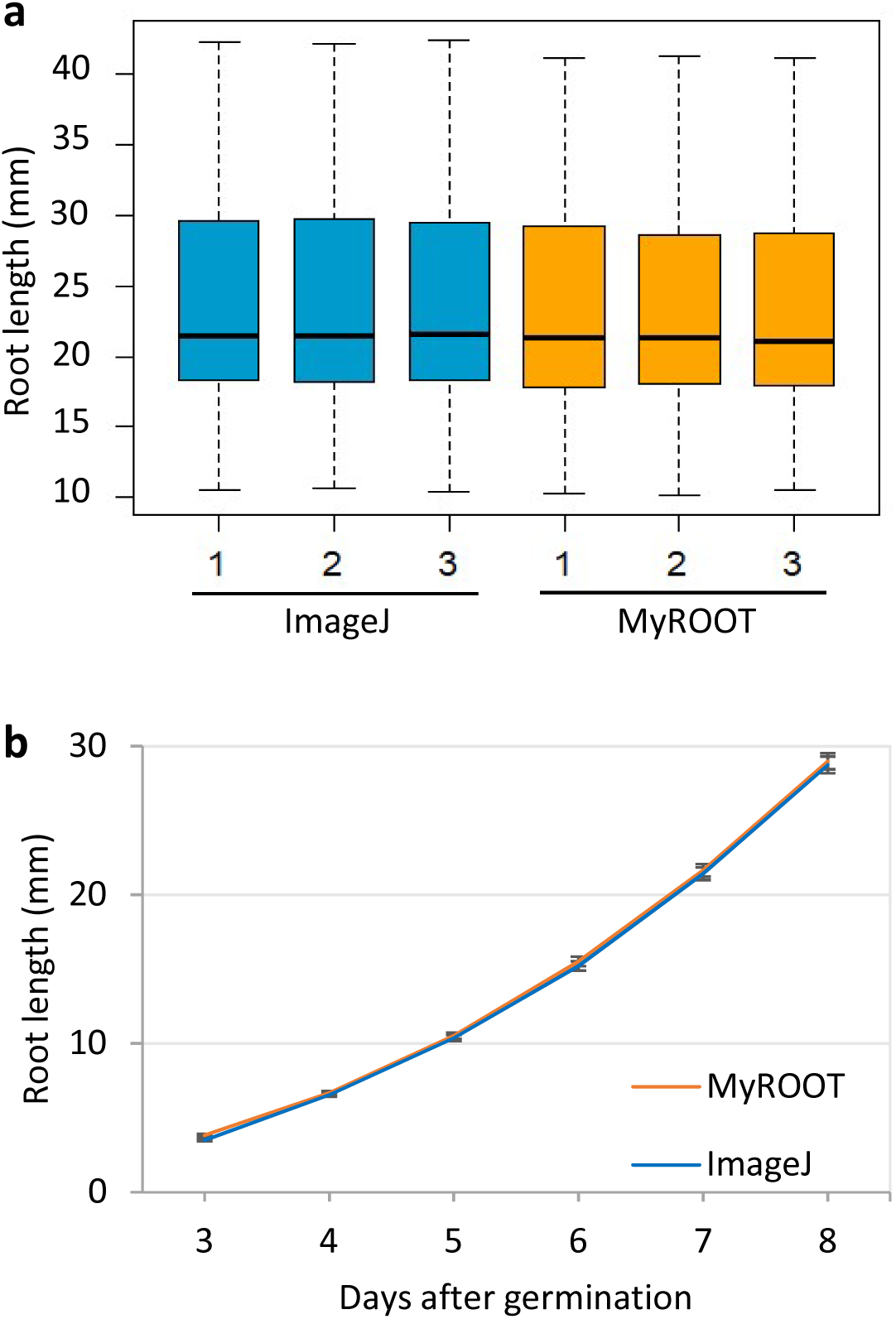
Validation of root length measurements. (**a**) Root length of 6-day-old Arabidopsis seedlings (3 plates, n=89). Measurements were performed by three different people using either the ImageJ tool or MyROOT. (**b**) Root length of Arabidopsis seedlings over 6 days (from 3 DAG to 8 DAG, n>116). Measurements were performed using either ImageJ or MyROOT. Error bars indicate the standard error. Seedlings that were not measured by MyROOT in at least 4 time points were discarded.

In addition, we also evaluated the time required by MyROOT to determine root lengths, and compared it to the time needed for manual measurements. Importantly, we found that our software significantly reduces the necessary time. MyROOT reduces the time required to measure one plate by approximately half (Fig. S2).

### Hypocotyl detection

One of the main advantages of MyROOT is its ability to identify the hypocotyls of the growing seedlings. This characteristic is important for accurately determining the start point of each root. The hypocotyl detection process is based on visual features extracted (appearance and color) from the image. These features were used to generate a hypocotyl model by introducing 1,259 hypocotyls of seedlings of different ages and characteristics and 7,915 samples with background information (see Material and Methods). The learned model is able to determine whether a given sample is a hypocotyl or not. To extract visual features, we implemented the histogram of oriented gradient (HOG) ^26^ method. The HOG method is based on the orientation of the contours in the image, and generates a histogram that represents the appearance/shape of the sample. For extracting color features, color distribution histograms representing the amount of color in a given sample area are used (Fig. 5a). To train the model, we implemented a linear support vector machine classifier that uses appearance and color features from the hypocotyl images. This classifier generates the best hyperplane that classifies samples as positive (hypocotyls) and negative (no hypocotyls) examples. During the hypocotyl detection stage, the sliding window approach ^27^ is used to perform an exhaustive search for hypocotyls. Finally, by keeping the highest scored windows as true positives, polynomial regression is used to define a curve that passes through all the detected hypocotyls. Although the user can manually insert the location of the hypocotyls, this curve enables the position of undetected hypocotyls to be estimated, and thus corrects the curve tracing. The intersection between the hypocotyl detection curve and each root is used to define the root start point.

**Figure 5.**
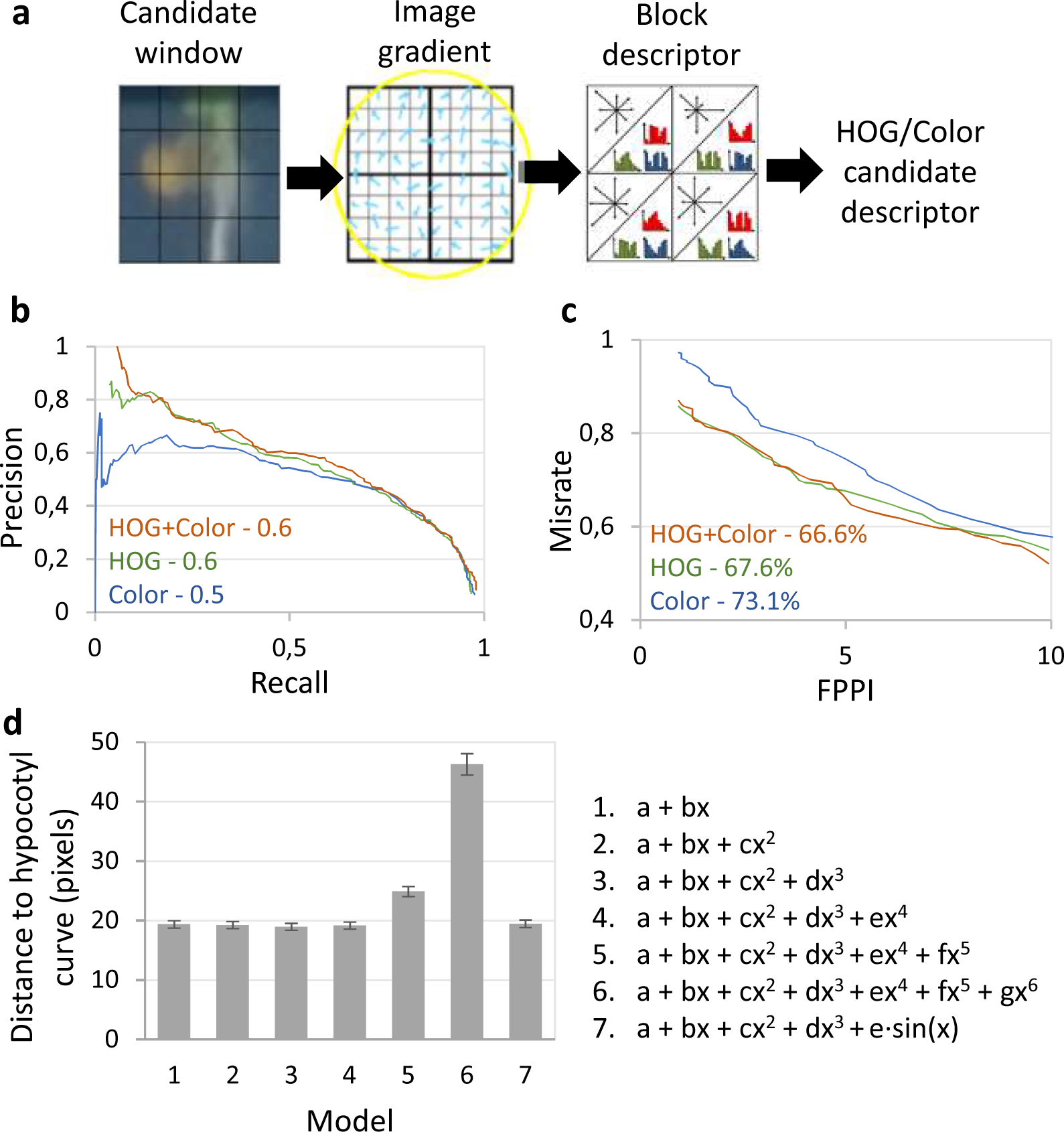
Hypocotyl detection method and validation. (**a**) Scheme of the hypocotyl detection method. A candidate window is defined as a square area inside the image. In order to describe a candidate, appearance/shape (HOG) and color information are extracted. Appearance information is extracted to calculate the gradient of the image (i.e., the direction of the contours within the image at each pixel). Histograms of Oriented Gradient (HOG) and the histograms of color are calculated over regular spaced, non-overlapping cells inside the candidate window (forming the block descriptor). Finally, all color/HOG cell histograms are concatenated to obtain the candidate window description. (**b**) Precision-Recall curve for three different models of hypocotyl detection (HOG, Color and HOG+Color). The curve is obtained by changing the threshold that defines the frontier between positive and negative samples. For each threshold, the precision (well classified ratio) and the recall (poor classified ratio) were calculated. The area under the curve represents the robustness of the classifier, with a higher value indicating greater robustness (a higher well classified ratio to poor classified ratio over the entire range of the classifier). (**c**) False Positives Per Image (FPPI) curve for three different models of hypocotyl detection (HOG, Color and HOG+Color). The curve plots the miss rate against the FPPI. In this way, the average miss rate over a specific FPPI range (1 to 10) represents the sensitivity of the classifier to not miss good samples and keep the false positive ratio low. (**d**) The average distance in pixels between the real hypocotyl position and the point of intersection between the root and the polynomial regression curves, for polynomial regression curves of orders 1 to 6 and an extra model including a sine component. Error bars indicate the standard error.

We first evaluated our hypocotyl detection process in terms of different hypocotyl detection models. Both the precision-recall curve (Fig. 5b) and the number of false positives per image (FPPI; Fig. 5c) were calculated for three different models that differ in the type of feature they use for describing hypocotyls: only color information, only appearance information (via HOG features), or both types of information (HOG + color).

Upon analyzing the precision-recall curve of each model, we found the HOG + color model to be the most robust (Fig. 5b). In the case of FPPI, the lowest miss rate was also found when using the HOG + color model (Fig. 5c). These results indicate that when considering both color and appearance (i.e., the HOG + color model), more hypocotyls are identified than when using only one of the features. Thus, this validates our MyROOT method because it incorporates both HOG and color information.

Next, we evaluated the influence of different regression curve models on the root measurement refinement used to set up the limits of individual roots (Fig. 5d). To create these curves, a regression upon the detected hypocotyls was performed. In order to define which regression model gives the better fit, we tested different polynomial models that were evaluated in terms of the average distance (in pixels) between the real hypocotyl position and the point of intersection between the root and the regression curve (Fig. 5d). The results indicate that when using a hypocotyl regression curve of order 4, a good balance between accuracy and flexibility that is able to account for small changes in hypocotyl position is reached. As such, we chose to employ this regression curve in our software.

## DISCUSSION

The advent of root imaging and phenotyping has aided in the understanding not only of plant organogenesis and development, but also of plant adaptation to changing environments ^19^ Here, we present MyROOT, a novel software for accurately measuring root length from images of Arabidopsis seedlings that are grown vertically in Petri dishes. In addition to its simple image acquisition method, this software minimizes user intervention through the automatic detection of the scale, root tips and hypocotyls. Compared to other available root software such as WinRhizo ^28^ or GROWSCREEN-Root^29^, which require specialized imaging equipment such as scanners or infrared light, MyROOT merely uses a standard digital camera.

One of the novel aspects of this software compared with other existing methods such as ImageJ and RootTrace ^23^ is that MyROOT has been trained to automatically identify hypocotyls, the shoot-root junction and the root tip of each seedling. Such automated identification is necessary for minimizing the manual intervention of the user and as such, the time dedicated to the root measurement task. In addition, MyROOT is also able to identify hypocotyls of different sizes and morphologies, an aspect that also makes it suitable for the phenotypic analysis of mutants or of plants grown in different conditions. Furthermore, in exceptional cases where the software might be unable to automatically recognize hypocotyls, they can actually be manually indicated by the user. In this way, we have designed MyROOT as a versatile tool that can be used in a wide range of developmental studies, including analyses of seedlings at different developmental stages and under different conditions.

MyROOT is a modularly designed software. It consists of a group of specialized algorithms able to detect and analyze the measuring tape, detect the roots, track the roots in a bottom-up fashion, and detect the hypocotyls. Therefore, any improvement to any of these components, or new algorithms for the determination of other features, can be easily included in subsequent versions of MyROOT. Examples of future improvements that could be included are the development of daily growth-monitoring algorithms that permit the detection of abnormal root growth patterns, the analysis of root system architecture beyond the primary root, and the identification of hypocotyls from other plant species. In the future, upgraded versions of our software could consist of a completely automatic operation connected to high-throughput facilities for massive root phenotyping.

## CONCLUSION

MyROOT is a software capable of semi-automatically measuring the length of the primary root of Arabidopsis seedlings. It automatically recognizes the scale of the image, and detects the hypocotyls and root tips from young seedlings growing vertically in agar plates. This information is then used to accurately calculate the root length of each individual plant. This software was designed in such a way that only a simple image of the plate is required for analysis. Importantly, MyROOT is even able to recognize hypocotyls of different ages and morphologies, and can thus be applied in a large range of experiments.

Here, our validation experiments demonstrate the high precision of measurements made with MyROOT, thereby proving that this software can be used within the research community to perform high-throughput experiments in a less time-consuming manner.

## MATERIAL AND METHODS

### Plant material and growth conditions

Wild type Col-0 *Arabidopsis thaliana* seeds were surface sterilized with a 5-min incubation in 1.5% sodium hypochlorite, followed by five washes in distilled sterile water. Seeds were stratified for 48 h at 4°C in the dark to synchronize germination. Seeds were sown in 12x12-cm plates containing ½ Murashige and Skoog (MS) medium without sucrose and supplemented with vitamins (0.5MS-). Seeds were distributed individually in the plate in two rows with around 15 seeds per row. The plates were incubated for 3 to 8 days vertically oriented under long day conditions (16 hours of light and 8 of hours dark) at 22°C and 60% relative humidity.

### Plant Imaging and computer settings

Images were taken with a D7000 Nikon camera. The pre-defined image acquisition conditions consist of placing the camera 50 cm above the plate with an illuminated support and the following settings: aperture 13, shutter speed 10, ISO 100 and Zoom x35. The plates were placed face down on a black surface and with a ruler (at least 1 cm long) horizontally positioned on top (Fig. S1). The images were saved in JPEG format (size between 2.5 and 2.7 MB per image).

MyROOT and ImageJ were run in a Intel^®^ Core^™^ i7-6700 CPU computer.

### Hypocotyl detection model

The software was trained to identify hypocotyls by using 1,259 positive examples (hypocotyls) and 7,915 background and negative examples (parts of the image that did not contain hypocotyls). The positive samples correspond to Col-0 wild type, *bril-116*, and a transgenic line overexpressing BRI1-GFP, which have morphologically different hypocotyls as shown in ^11^.

### Algorithms

MyROOT has been developed in Matlab (version 8.3.0.532. Natick, Massachusetts: The MathWorks Inc., 2014). It will be made available to the plant sciences community through the Plant Image Analysis website (plant-image-analysis.org; ^30^) a standalone executable application.

## ACKNOWLEDGMENTS

We would like to thank Caño-Delgado Lab members for helping with the manual root length measurements and comments on the manuscript. A.I.C-D. is a recipient of a BIO2016-78955 grants from the Spanish Ministry of Economy and Competitiveness and a European Research Council, ERC Consolidator Grant (ERC-2015-CoG – 683163). I.B-P. is funded by the FPU15/02822 grant from the Spanish Ministry of Education, Culture and Sport. D.B-E. is contracted with the BIO2016-78955 grant in the A.I.C-D laboratory. CRAG is funded by “Severo Ochoa Programme” from Centers of Excellence in R&D 2016-2019 (SEV-2015-485 0533).

## AUTHOR CONTRIBUTIONS

A.I.C-D. conceived the idea. A.G. and X.S. developed the algorithms for the method. A.G., X.S., I.B-P. and D.B-E performed the validation experiments. I.B-P. and D.B-E. acquired the dataset. X.S. and A.I.C-D. designed and supervised the study. I.B-P., A.G. X.S. and A.I.C-D. wrote the manuscript.

## FIGURE LEGENDS

**Supplementary Figure 1.** Laboratory setup for taking pictures of the plates. The image shows the position of the lights, the camera and the plate to be analyzed, all positioned over a black surface.

**Supplementary Figure 2.** Evaluation of the time required to measure root length. Time (in seconds) required for three different scientists to measure the root length of two different plates containing one and two rows of seedlings respectively. The measurements were done with MyROOT and ImageJ. Error bars indicate the standard error.

#### BOX 1: Installation guide for MyROOT software

1. Execute MyAppInstaller_mcr.exe.
2. In the Root Analysis Installer window press the *Next* button.
3. In the Installation Options window select the folder where you wish to install the software. By default, the selected folder is C:\Program Files\La Salle – Universidad Ramon Llull\Root_Analysis.
4. Mark the option *Add a shortcut to the desktop*.
5. Press the *Next* button.
6. In the Required Software window select the installation folder for the MATLAB Runtime (by default C:\Program Files\MATLAB\MATLAB Runtime)
7. Press the *Next* button.
8. In the License Agreement window mark the option *Yes*.
9. Press the *Next* button.
10. In the Confirmation window press the *Install* button.
11. Copy the HypocotylDetection folder to the Desktop (this folder is in the same folder as the ․exe file used for the installation).
12. Open the software by pressing in the MyROOT Desktop icon.

#### BOX 2: Brief user guide for MyROOT software

1. Open the software in a PC.
2. Select and load the image to process by pressing the *LOAD IMG* button. Enter an image resize factor between 0 and 1 in the Scale edit box to reduce the size of the image and speed up the processing of high-resolution images.
3. Obtain the pixels-to-millimeters scale factor by pressing the *Ruler ID* button. If needed, edit the Ruler Threshold value to modify the sensitivity of the ruler detector and repeat step 3.
4. Start the root segmentation process by pressing the *Root Mask* button. Select the area where the roots are present by clicking in the image. Double click in one of the vertex to start generating the mask. In case the result is not satisfactory (e.g., over-segmented roots), modify the sensitivity factor in the Root Threshold edit box and repeat step 4.
5. Enter a value in the Root Length Threshold box to indicate the minimum percentage with respect to the longest root to be measured.
6. Start the root tracking and measurement process by pressing the *Root Detection* button.
7. Enter the path of the files containing the pre-trained hypocotyl detection models in the Hypocotyl Model Path edit box.
8. Optionally, to perform hypocotyl detection based on color descriptors only, check the *Only Color* checkbox, and to conduct a channel-wise color normalization process check the *Norm Color* checkbox.
9. Optionally, modify the threshold of the linSVM classifier by modifying the value in the edit box located next to the Hypocotyl Detection button.
10. Start the hypocotyl detection process by pressing the *Hypocotyl Detection* button.
11. If some of the hypocotyls were undetected, insert them manually by using the *Add* button. Press the *Enter* key and press the *Root Refinement* button to update root length measurements.
12. Remove the undesired roots from the measurement by typing the root identifier in the ID edit box and pressing the *Remove* button. Press the *Visualize* button to refresh the image presented on MyROOT’s visualization canvas.
13. Enter the path where you would like the results to be stored in the Results Path edit box.
14. Choose the type of data you want to save by checking the corresponding checkboxes.
15. Optionally, type an identification suffix that will be appended to the stored file names via the Root Label edit box.
16. Save the results by pressing the *SAVE* button.

## REFERENCES

1 Kuijken, R. C., van Eeuwijk, F. A., Marcelis, L. F. & Bouwmeester, H. J. Root phenotyping: from component trait in the lab to breeding. Journal of experimental botany 66, 5389–5401, doi:10.1093/jxb/erv239 (2015).

2 Dolan, L. et al. Cellular organisation of the Arabidopsis thaliana root. Development 119, 71–84 (1993).

3 Ishikawa, H. & Evans, M. L. Specialized zones of development in roots. Plant physiology 109, 725–727 (1995).

4 Iyer-Pascuzzi, A., Simpson, J., Herrera-Estrella, L. & Benfey, P. N. Functional genomics of root growth and development in Arabidopsis. Current opinion in plant biology 12, 165–171, doi:10.1016/j.pbi.2008.11.002 (2009).

5 Beemster, G. T. & Baskin, T. I. Analysis of cell division and elongation underlying the developmental acceleration of root growth in Arabidopsis thaliana. Plant physiology 116, 1515–1526 (1998).

6 Verbelen, J. P., De Cnodder, T., Le, J., Vissenberg, K. & Baluska, F. The Root Apex of Arabidopsis thaliana Consists of Four Distinct Zones of Growth Activities: Meristematic Zone, Transition Zone, Fast Elongation Zone and Growth Terminating Zone. Plant signaling & behavior 1, 296–304 (2006).

7 Takatsuka, H. & Umeda, M. Hormonal control of cell division and elongation along differentiation trajectories in roots. Journal of experimental botany 65, 2633–2643, doi:10.1093/jxb/ert485 (2014).

8 van den Berg, C., Willemsen, V., Hendriks, G., Weisbeek, P. & Scheres, B. Short-range control of cell differentiation in the Arabidopsis root meristem. Nature 390, 287–289, doi:10.1038/36856 (1997).

9 Jürgens, G., Mayer, U., Busch, M., Lukowitz, W. & Laux, T. Pattern formation in the Arabidopsis embryo: a genetic perspective. Philosophical transactions of the Royal Society of London. Series B, Biological sciences 350, 7, doi:10.1098/rstb.1995.0132 (1995).

10 Benfey, P. N. et al. Root development in Arabidopsis: four mutants with dramatically altered root morphogenesis. Development 119, 57–70 (1993).

11 Gonzalez-Garcia, M. P. et al. Brassinosteroids control meristem size by promoting cell cycle progression in Arabidopsis roots. Development 138, 849–859, doi:10.1242/dev.057331 (2011).

12 Potuschak, T. et al. EIN3-dependent regulation of plant ethylene hormone signaling by two arabidopsis F box proteins: EBF1 and EBF2. Cell 115, 679–689 (2003).

13 Hauser, M. T., Morikami, A. & Benfey, P. N. Conditional root expansion mutants of Arabidopsis. Development 121, 1237–1252 (1995).

14 Cano-Delgado, A. I., Metzlaff, K. & Bevan, M. W. The eli1 mutation reveals a link between cell expansion and secondary cell wall formation in Arabidopsis thaliana. Development 127, 3395–3405 (2000).

15 Mouchel, C. F., Briggs, G. C. & Hardtke, C. S. Natural genetic variation in Arabidopsis identifies BREVIS RADIX, a novel regulator of cell proliferation and elongation in the root. Genes & development 18, 700–714, doi:10.1101/gad.1187704 (2004).

16 Ubeda-Tomas, S. et al. Root growth in Arabidopsis requires gibberellin/DELLA signalling in the endodermis. Nature cell biology 10, 625–628, doi:10.1038/ncb1726 (2008).

17 Gonzalez-Garcia, M. P. et al. Single-cell telomere-length quantification couples telomere length to meristem activity and stem cell development in Arabidopsis. Cell reports 11, 977–989, doi:10.1016/j.celrep.2015.04.013 (2015).

18 Pfister, A. et al. A receptor-like kinase mutant with absent endodermal diffusion barrier displays selective nutrient homeostasis defects. eLife 3, e03115, doi:10.7554/eLife.03115 (2014).

19 Lobet, G. Image Analysis in Plant Sciences: Publish Then Perish. Trends in plant science 22, 559–566, doi:10.1016/j.tplants.2017.05.002 (2017).

20 Yazdanbakhsh, N. & Fisahn, J. High-throughput phenotyping of root growth dynamics. Methods Mol Biol 918, 21–40, doi:10.1007/978-1-61779-995-2_3 (2012).

21 Slovak, R. et al. A Scalable Open-Source Pipeline for Large-Scale Root Phenotyping of Arabidopsis. The Plant cell 26, 2390–2403, doi:10.1105/tpc.114.124032 (2014).

22 Clark, R. T. et al. High-throughput two-dimensional root system phenotyping platform facilitates genetic analysis of root growth and development. Plant, cell & environment 36, 454–466, doi:10.1111/j.1365-3040.2012.02587.x (2013).

23 French, A., Ubeda-Tomas, S., Holman, T. J., Bennett, M. J. & Pridmore, T. High-throughput quantification of root growth using a novel image-analysis tool. Plant physiology 150, 1784–1795, doi:10.1104/pp.109.140558 (2009).

24 Remmler, L., Clairmont, L., Rolland-Lagan, A. G. & Guinel, F. C. Standardized mapping of nodulation patterns in legume roots. The New phytologist 202, 1083–1094, doi:10.1111/nph.12712 (2014).

25 Otsu, N. A Threshold Selection Method from Gray-Level Histograms. IEEE Transactions on Systems, Man, and Cybernetics 9, 5, doi:10.1109/tsmc.1979.4310076 (1979).

26 Dalal, N. & Triggs, B. Histograms of Oriented Gradients for Human Detection. Computer Society Conference on Computer Vision and Pattern Recognition 1, 8, doi:10.1109/CVPR.2005.177 (2005).

27 Glumov. Detection of objects on the image using a sliding window mode. Optics & Laser Technology 27, 9 (1995).

28 Arsenault, J., Poulcur, S., Messier, C. & Guay, R. Winrhizo: A root measuring system with a unique overlap correction method. HortScience 30 (1995).

29 Nagel, K. et al. Temperature responses of roots: impact on growth, root system architecture and implications for phenotyping. Functional Plant Biology 36, 12 (2009).

30 Lobet, G., Draye, X. & Perilleux, C. An online database for plant image analysis software tools. Plant methods 9, 38, doi:10.1186/1746-4811-9-38 (2013).

